# Magnetic bioprinting of stem cell-based tissues

**DOI:** 10.1101/2022.12.23.521759

**Authors:** Aurore Van de Walle, Jose Efrain Perez, Claire Wilhelm

## Abstract

An attractive approach in cell therapies and medically oriented nanotechnologies is to interface magnetic nanoparticles with cells. This will supply the cells with sufficient magnetization for theranostic applications and for external magnetic field manipulation.

In tissue engineering, one challenge is to produce tissue analogues that are large, precisely organized, and responsive to stimuli, preferably without the need for an artificial supporting scaffold. One powerful tool for such biofabrication is certainly the bioprinting technology.

In magnetic tissue engineering, it appears possible to use magnetic forces to manipulate cells, both individually and within aggregates, and thereby to produce three-dimensional artificial tissues with inherent capacities for further physical stimulation, a possibility that bioprinting does not offer yet.

We here introduce the feasibility of using magnetic forces created by external (micro)magnets to form 3D tissue-like scaffold-free structures. Because stem cells are essential in tissue engineering, such magnetic technologies were developed with magnetized stem cells, and applied for instance to vascular or cartilage tissue engineering. One precondition to this approach, which lies in the magnetization of (stem) cells endowed through internalization of iron oxide magnetic nanoparticles, is to ensure the safety of magnetic nanoparticles with respect to cellular functions, which is initially discussed.

Finally, we introduce a magnetic tissue stretcher which, in a single step, allows to create a tissue composed of any type of component cell, then to mature it, stimulate it by compression or stretching at any desired frequency, e.g. cyclically, opening new possibilities in the cardiac muscle tissue engineering field.

## Introduction

Recent developments in tissue engineering have focused on techniques yielding spatial organization of cells. Tissues are hierarchical structures whose proper functioning depends on their precise, complex architecture. This framework will need to be reproduced *in vitro* if we are to succeed in creating tissues with the same functions as their *in vivo* equivalents. Initial cell seeding approaches generally lacked control of cell organization[1]. To elicit spatial control, cell adhesion guidance has been tested by different strategies[2]. Cells can be (i) passively patterned on engineered substrates (e.g. nano-structuration[3] or biochemical modifications[4–8]) with the risk of irreversibly affecting the cell adhesion properties, or (ii) actively patterned by the remote application of a force (e.g. electric[9,10] or optic[11,12]). To avoid permanent cell adhesion, reversible substrates were proposed[13], yet they are expensive and time-consuming.

Active patterning of cells to build structural tissues can be achieved by 3D bioprinting. It traditionally refers to the use of a printer with a three-axis mechanical platform moving accordingly to defined coordinates that controls the deposition of biomaterials, either encapsulating cells or loaded with them afterward. Bioprinting methodologies are still evolving, with new bioinks tailored to the tissue specificities or made of cells only[14,15], and with higher resolution printers that allow multi-material deposition for the assembly of more complex structures[16–18] while becoming cost-effective[19]. Teams succeeded in producing human-scale correctly-shaped skin, bone, cartilage, and skeletal muscle[20–22]. Some studies even moved to 4D, with materials able to change shape[23], and combined such materials with anisotropic particles that can be controlled magnetically to generate unprecedented microstructural features[24].

The 3D bioprinting of tissues can also be performed magnetically. In this case, cells are organized into specific 3D patterns using external magnetic forces. A pre-requirement to this remote organization of cells is their labeling with magnetic nanoparticles to render them magnetic. It should be noted that magnetic nanoparticles have already made their way into a variety of medical applications, including magnetic resonance imaging or iron replacement therapy[25]. Their use is also assessed for thermal cancer therapy such as hyperthermia[26,27] or targeting and drug delivery[28,29]. Consequently, they are considered as good candidates for such applications, provided the control of their design and internalization dose[30].

The magnetic bioprinting strategy has been initially explored to magnetically engineer cell sheets[31–33], taking advantage of cell magnetization to form and organize tissue constructs in the presence of a magnetic field gradient that resulted in successive cell layering. The magnetic formation of cell spheroids has also been achieved. For instance, it has been performed by magnetic levitation that consists in the application of an external magnetic to levitate cells at the air-medium interface of a culture and results in cohesive 3D cell assembly[34,35]. Another strategy is to use magnetic forces to create thick 3D tissue-like structures[36,37]. Magnetic patterning of cells has been successful in building submillimetric scale-tissues, as well as to promote cell assembly with tuneable size, controlled geometry, and without the need of a supporting matrix[38–42]. Relative to other patterning techniques, magnetic forces provide access to the third dimension thanks to the long range of magnetic interactions. In addition, they are totally nonspecific to cell functions and the forces applied can be precisely controlled.

Originally, these magnetic bioprinting approaches to tissue engineering and manipulation were introduced for tissue formation, in which magnetic forces are used to organize single cells within a 3D construct. It has now moved to 4D magnetic bioprinting with the stimulation of the 3D engineered constructs by remote physical forces. The rationale is then to magnetically arrange millions of cells into a 3D construct and exploit the magnetic properties of the construct to stimulate it mechanically[43].

Tissue engineering strategies are generally grounded on stem cells that are a potentially unlimited source of tissue due to their proliferation and differentiation capacities. They can generate one or several specialized cell types depending on their level of potency. Embryonic stem cells (ESCs) are pluripotent and can develop into any cell type, except those of the placenta[44]. They are found at an early embryonic stage and give rise to the different organs during ontogenesis. Although less numerous than in embryos, stem cells are also present in adults, where they ensure maintenance and in some cases the regeneration of tissue. Among them, the multipotent mesenchymal stem cells (MSCs) have the capacity to differentiate into cell types derived from the mesenchyme, including osteoblasts, chondrocytes, adipocytes, and myocytes[45]. Endothelial progenitor cells (EPCs) are even more specialized, being integrated in the formation of new vessels and able to differentiate into mature endothelial cells[46].

Molecular determinants of stem cell differentiation pathways have now been identified, and it has been revealed that molecular signals are not the only factors involved in differentiation processes. Some physical features of the cellular microenvironment and its dynamic behavior also play a determining role[47–49]. For instance, the relative rigidity of a substrate can orient the differentiation of mesenchymal stem cells (MSCs) without the need for specific growth factors[50]. Thus, a soft substrate with a rigidity similar to that of the brain (1 kPa) orients MSC differentiation into neuronal cells, while a hard substrate comparable to bone tissue (100 kPa) gives rise to bone cells. In addition to matrix stiffness, the spatial organization of substrate molecules and cells can affect differentiation. For instance, the nanoscale patterning of cell-adhesion ligands on a substrate affects the spreading area of MSCs and their differentiation[51], and has been heavily investigated in the promotion of osteogenesis[52]. Another example is the necessity of cell-cell interactions for the initiation of chondrogenesis, which can be promoted by the confinement of MSCs in a 3D aggregate composed of several thousand cells[53]. The influence of dynamic external constraints such as fluid flow (shear stress) or compression has also been demonstrated, and the importance of specific and precise mechanical microstimulations is still being revealed. Indeed, it was shown that the capacity of EPCs to build new functional vessels is influenced by the shear stress generated by blood flow[54]. Flow conditions also encourage MSCs to differentiate into endothelial cells[55] and osteoblasts[56]. Mechanical loading, such as compression of 3D gels incorporating stem cells (generally MSCs), induces differentiation and strongly enhances the regeneration of bone and cartilage tissues by activating the appropriate mechanotransduction pathway[57–61]. Multiple biomaterial-based niches have been proposed in order to better understand the role of mechanical transduction in stem cell differentiation and to further develop the field of tissue engineering and regenerative medicine[62]. Matrix stiffness has for example been coupled with molecular factors to induce smooth muscle cell differentiation[63,64], and focus has been placed on trying to understand the molecular mechanisms[65] and subsequent gene response[66]. However, in these studies the constraint was applied globally, meaning that the cells could not be assembled into a particular architecture. The magnetic bioprinting strategy provides an option to organize cells in 3D and stimulate them remotely at the same time through the use of magnetic forces.

This review reports different works using the concept of magnetically engineered tissues based on stem cells whose differentiation is directed. Fabrication of vessels *de novo* can for example be achieved by organizing two different stem cell types (endothelial progenitors EPCs and MSCs)[67]. The formation of cartilaginous tissue can be accomplished via an initial magnetic aggregation of human MSCs followed by their differentiation along the chondrogenic pathway and further tissue building[36]. Magnetically induced cyclic stretching and compression of mouse ESCs can be performed to induce cardiomyogenesis[43]. The magnetic labeling of stem cells with magnetic nanoparticles will first be discussed, followed by the presentation of the principles of magnetic cell assembling and the dynamic behavior of the MSC magnetic aggregates.

### 1. Stem cells magnetization upon internalization of magnetic nanoparticles

#### 1.1. Cell uptake of nanoparticles: quantification by single-cell magnetophoresis

As most nanomaterials, magnetic nanoparticles are generally captured by the cells through the endocytosis pathway and concentrated within lysosomes (Figure 1A). One of the most used magnetic nanoparticles in the biomedical field are synthesized by co-precipitation of iron salts and stabilized by citrate adsorption on their surface to ensure colloidal stability via negative electrostatic repulsions. Figure 1B and 1C show transmission electron microscopy (TEM) images of MSCs and endothelial progenitor cells upon internalization of these types of nanoparticles, which appear densely confined within endosomes. Cells having incorporated the nanoparticles thus become magnetic: they can be attracted by any magnet creating a magnetic field gradient. Single-cell magnetophoresis relies precisely on this concept of magnetic migration to measure the magnetic moment of cells, or equivalently the amount of nanoparticles per cell (expressed usually in pg of iron)[68]. Figure 1D shows a sequence of magnetophoresis performed on magnetic MSCs. It also clearly demonstrates that the nanoparticles labeling provides the cells with sufficient magnetization to be manipulated by external magnets. The magnetophoretic quantification of the mass of iron per cell indicates the nanoparticle uptake. Figure 1E shows this uptake as a function of the dose of nanoparticles for MSCs, EPCs, and ESCs upon incubation with the nanoparticles for only 30 min. The nanoparticles presence within the cells can then be confirmed via direct observation after histological iron staining (Figure 1F).

**Figure 1:**
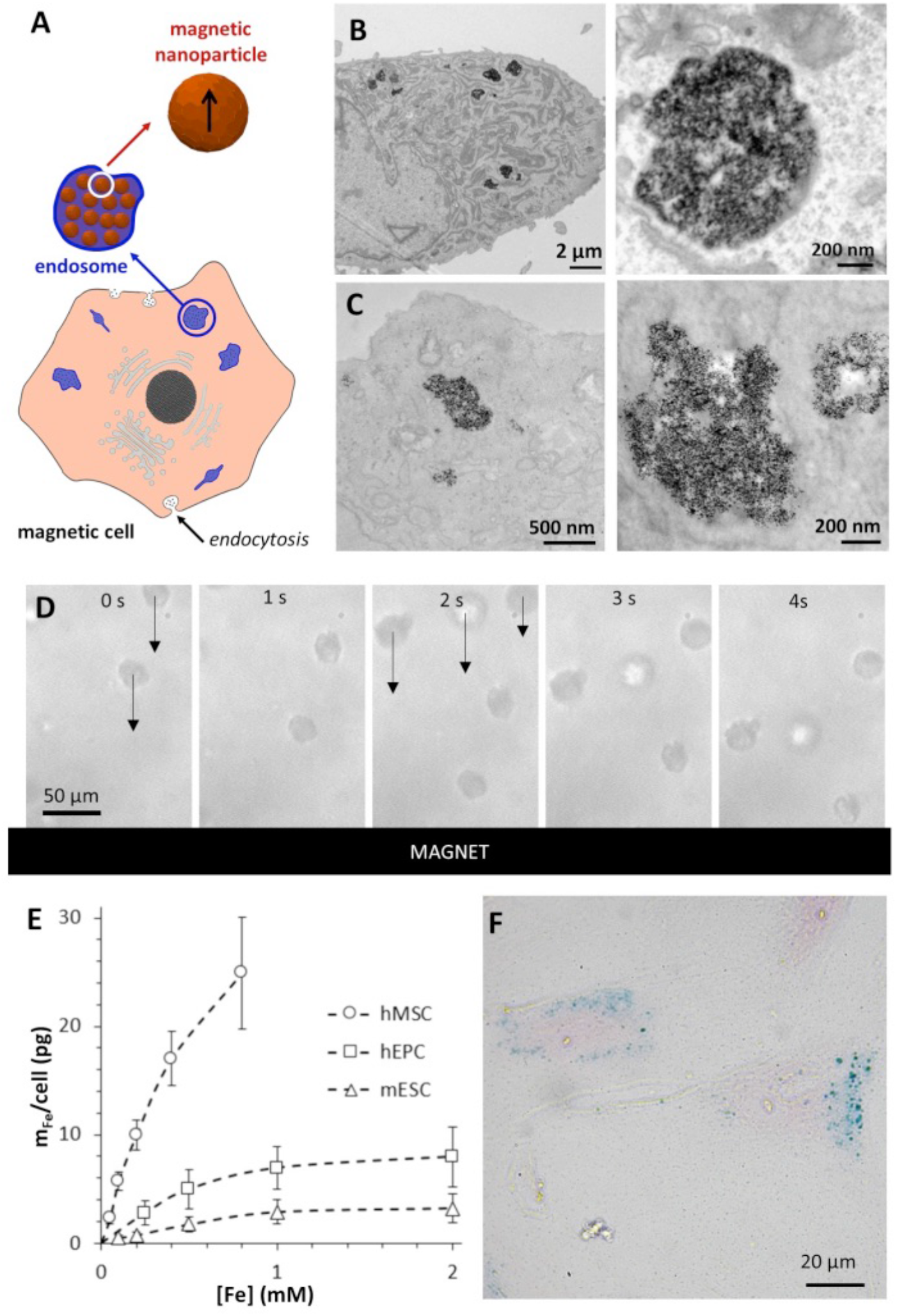
A. Schematic representation of magnetic nanoparticles endocytosis. B,C. Representative TEM images of EPCs (B) and MSCs (C) showing the nanoparticles in the endocytic compartments. D. Magnetophoresis sequence showing that labeled cells are attracted towards a magnet. Each cell is exposed to a magnetic force m_cell_ x gradB, where m_cell_ is the magnetic moment of the cell and gradB = 17T/m for the magnet used. The cell migrates towards the magnet at a constant velocity v_cell_ depending on the equilibrium between the magnetic force and the Stokes viscous force (6πηR_cell_v_cell_, with η being the water viscosity and R_cell_ being the cell radius). Here, the cells (MSCs) have a mean diameter of 24 ± 2 μm and a velocity of 46 ± 11 μm/s, meaning a calculated moment of 6 ± 2 × 10^−13^ A.m^2^, corresponding to a mass of iron per cell of 10 ± 3 pg, or about 10 million nanoparticless per cell. E. Uptake curves of MSCs, EPCs, and ESCs. The variations in internalization mass are essentially due to the cell diameter: 24 μm for the MSCs, 14 μm for the EPCs, (diameters when in suspension). F. Prussian blue staining of labeled MSCs showing the iron internalized into the intracellular compartments in blue and the viable cells in pink (counterstained with nuclear fast red).

#### 1.2. Impact of nanoparticles internalization on stem cell function

For therapeutic and diagnostic applications of magnetically labeled cells, it is crucial to first verify the biological safety of intracellular magnetic nanoparticles. The labeling method (with nanoparticles made by co-precipitation, in aqueous medium, with or without polymer coating) has already been tested on many cell types from different species, in primary and secondary cultured cells, immune cells, malignant cells and muscle cells, among others. The labeling did not affect cell viability, proliferation or the therapeutic and functional properties of the tested cell types, neither *in vitro* or *in vivo*[69–71]. These magnetic nanoparticles are thus generally considered biocompatible.

By contrast, the question of differentiation capacity after magnetic labeling has been debated in the context of MSC differentiation. Using a variety of magnetic nanoparticles with an iron oxide core, no alteration but rather a positive effect in the differentiation capacity of human MSCs was demonstrated by different groups concerning adipogenic and osteogenic differentiation[72,68,73–82,30,83]. Oppositely, the chondrogenic differentiation has been discussed controversially, with some studies showing no effects[73–76,84], whereas others demonstrated inhibition and even failure of chondrogenesis[77,78,85].

Using the nanoparticles produced by co-precipitation, it was demonstrated that the differentiation of MSCs was influenced by magnetic nanoparticle aggregation and by the amount of intracellular iron, with this iron “dose” likely being an important determinant of nanoparticle toxicity[30,86]. The amount of iron internalized by each cell depends mainly on the external iron concentration (or nanoparticle density) and on the incubation time. For MSCs, the iron uptake saturates at about 30 pg, with higher iron values obtained being explained by nanoparticle aggregation on the outer surface of the cells[87]. Regardless of the intracellular amount and aggregation configuration of iron oxide nanoparticles, magnetic labeling did not affect the survival or proliferation of MSCs. Moreover, the osteogenic and adipogenic differentiation pathways were neither modified, regardless of the magnetic labeling conditions. In contrast, chondrogenesis was inhibited by high intracellular iron content and by nanoparticle aggregation on the outer cell surface[87].

The differentiation capacities of ESCs were rarely documented after magnetic labeling. Au et al.[88] demonstrated that magnetic nanoparticles associated with polylysine do not impact cardiac differentiation. Similarly, labeling of ESCs with magnetic nanoparticles wrapped with stearic acid-grafted PEI600 did not impact the expression of ectoderm, endoderm, and mesoderm markers[89]. In our work[43], magnetic labeling of mouse ESCs was achieved with anionic magnetic nanoparticles and the absence of a negative impact on their differentiation towards the mesoderm pathway was demonstrated.

Concerning EPCs, the magnetic labeling had no negative impact on the cell phenotype, in particular their viability, proliferation and migration capacities, as well as vascular tube formation in Matrigel[90].

### 2. 3D magnetic patterning with magnetically labeled stem cells: an alternative to bioprinting

Stem cells, via a simple incubation with magnetic nanoparticles, can thus be labeled with magnetic nanoparticles and guided at a distance using magnetic forces. Magnetic cell patterning can in this way be achieved within a supporting matrix, or without, by the formation of cellular aggregates compacting as spheroids that can be fused to form larger tissues. This magnetic fusion of spheroids to form large structures is quite similar to bioprinting techniques that rely on spheroids as building blocks for tissue modeling[91].

#### 2.1. Development of local micro-magnetic devices for controlling cellular organization

The aim is therefore to modulate, in space and time, the way in which cells assemble by applying a local magnetic constraint. This “magnetic patterning” takes place in 3D and allows controlling the geometric pattern of cells and their compactness[38].

Soft-iron tips with a diameter of 0.8 mm, or plates with a thickness of 0.5 mm, can be used to generate magnetic “traps” with cylindrical or flat symmetries by applying an external permanent magnet (Figure 2). Hollow cylindrical magnets can also be used to create tubular patterns.

**Figure 2:**
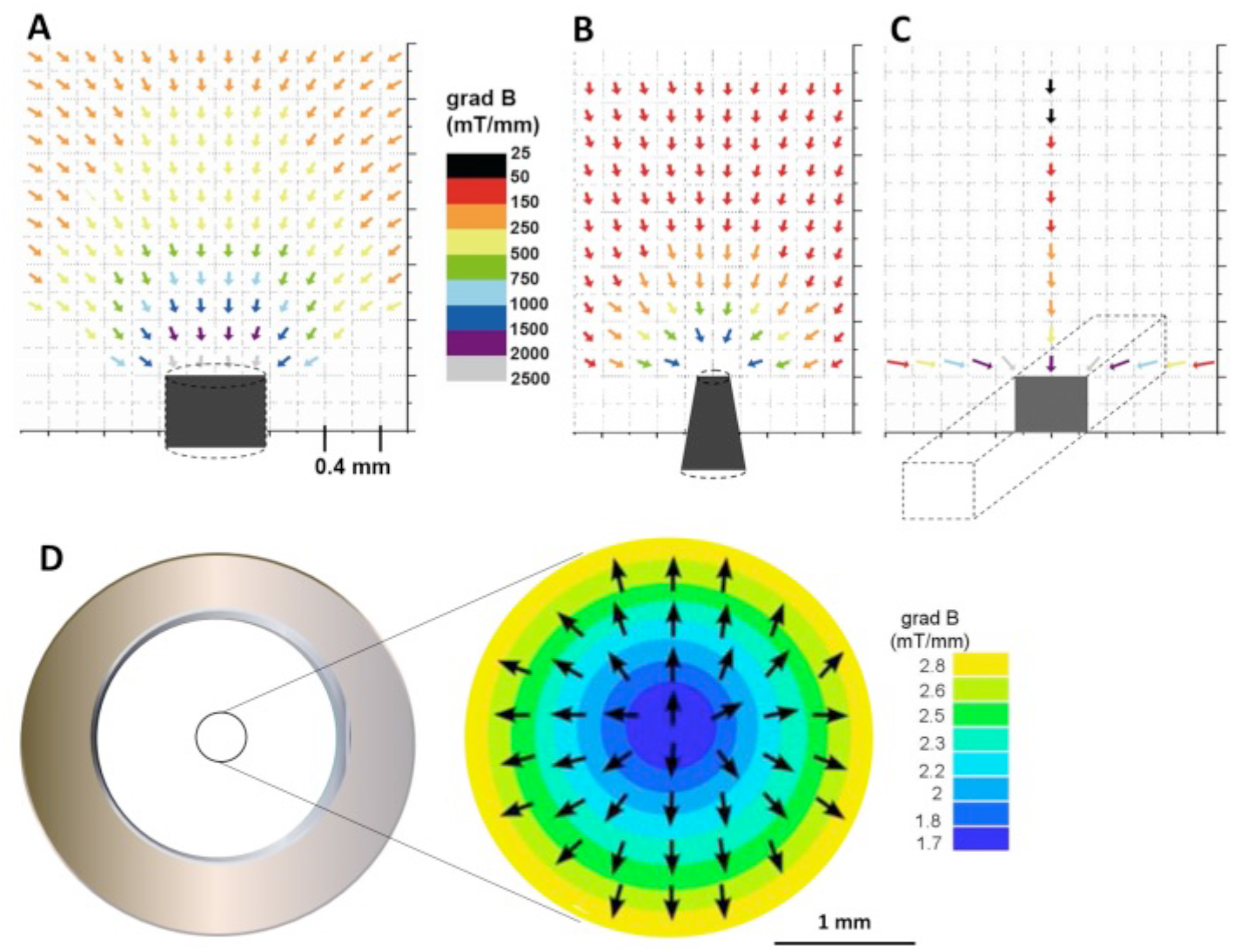
Map of the field gradients generated by a cylindrical tip (A), a truncated tip (B), a line (C), all made of soft iron, and (D) map in the center of a hollow disk magnet.

The field gradients are in the range 2-100 mT/mm, which is sufficient to attract suspended cells on millimetric scales (force exerted on a cell labeled with 10 pg of iron: 1-60 pN). It is then possible to form spherical or linear aggregates of thousands or hundreds of thousands of cells.

#### 2.2. Magnetic formation of cellular aggregates: a rapid way to probe cell-to-cell interactions

Cellular spheroids have emerged as alternative *in vitro* models to 2D cultures as they integrate the 3D geometry of the physiological environment. These spheroids can for instance be used to screen drugs[92], and a first necessary step for their formation is the gathering of cells in close contact to each other. The immediate application of the 3D magnetic patterning is to maintain, for several hours or days, a large number of cells in a well-defined region of space by transient application of an external magnetic field. The magnetic field gradients generated by the soft iron tips shown in Figure 2A & 2B are sufficiently powerful to entrap labeled cells. The advantages of this type of magnetic confinement are that it does not require prior chemical surface treatment, and it can yield 3D cell aggregation almost instantaneously. Magnetic molding can lead to the obtention of cohesive tumor spheroids with defined diameter within 24h[93]. It is a unique way to put cells in contact in a 3D environment, and remarkably, to explore the establishment of cell-to-cell liaisons during the early stages of tissue formation. Figure 3 shows the magnetic assemblies obtained over a magnetic tip for multiple cell lines, including human MSCs. The tip is maintained for 10 min, then removed, and the spontaneous dynamic behavior of the cell aggregates upon the release of the magnetic force directly mirrors the tissue cohesion and reflects cell-to-cell adhesion properties.

**Figure 3:**
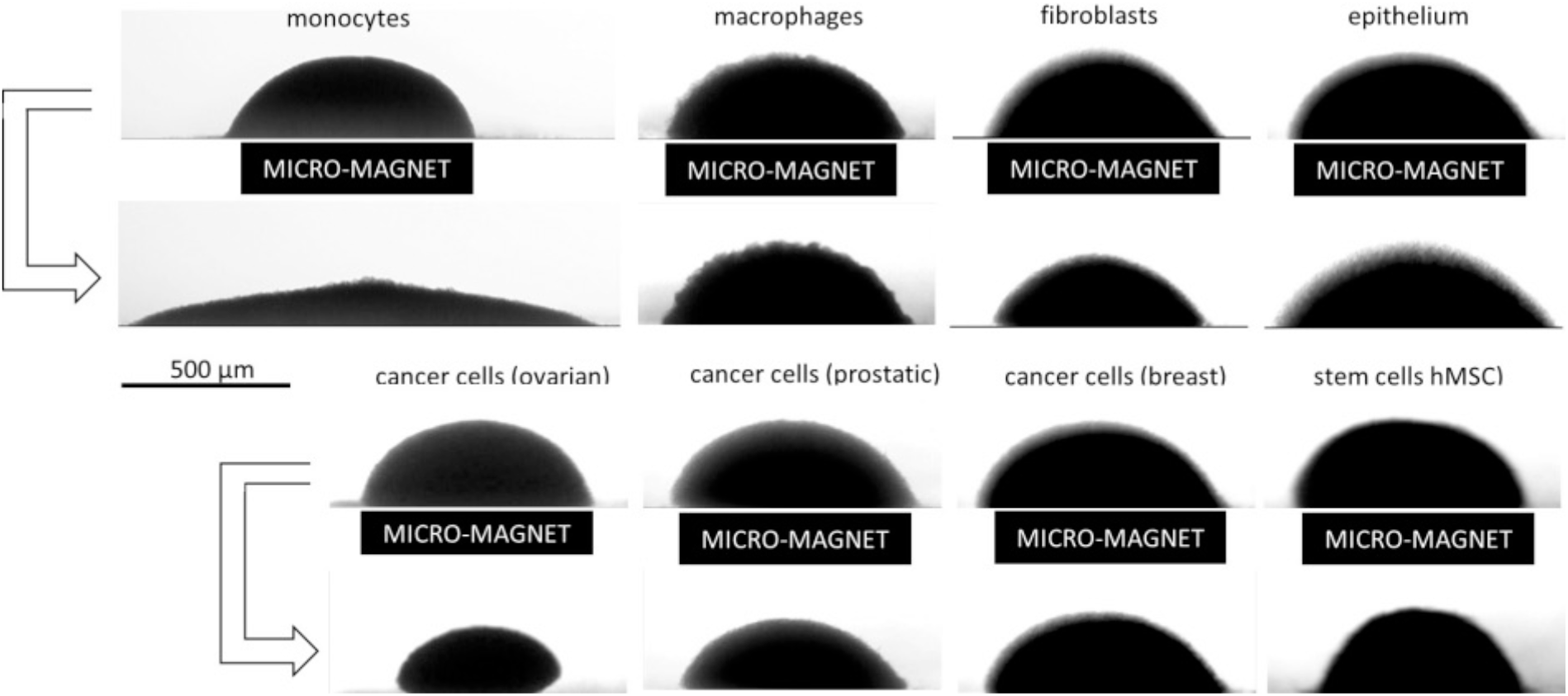
Formation of cell aggregates at the submillimetric scale by application of local intense field gradients generated beneath the substrate by magnetic tips. The magnet is maintained for 10 minutes before being removed. The second and fourth rows of images show aggregate evolution after 2 hours of spontaneous relaxation. The aim is to study the cellular dynamics during the first moments of tissue development. We observe a behavior that varies from granular (similar to a heap of sand that flows when the constraint is removed) to viscoelastic, with the most cohesive aggregates being based on cancerous and stem cells.

Understanding how cells interact to form functional structures is a major challenge. Besides, a precise knowledge of cellular behavior within tissues is necessary to meet the goals of tissue engineering and regenerative medicine.

In recent decades, driven by the differential adhesion hypothesis advanced by Steinberg in the 1960s, significant progress has been made in the description and modeling of tissue morphogenesis. A tissue aggregate is now described most of the time as a liquid droplet, defined by a surface tension that reflects the internal cellular adhesions. However, all existing experiments apply solely to aggregates already formed from cells exhibiting strong intercellular interactions (embryonic, heart, liver, retina cells, etc.). In contrast, the initial steps leading to the formation of a tissue structure, through cell-cell adhesion, have so far been described only for small numbers of interacting cells (mostly simple doublets).

Thanks to the magnetic cell aggregation technique allowing the assembly of millions of cells instantaneously, it has become possible to probe the establishment of 3D cell-cell adhesion at early stages within cell aggregates. By testing a large number of cell types, it was clearly evidenced that cell aggregates behave like complex materials, with a transition from wet granules to contractile structures, controlled by cell-cell interactions[39]. Importantly, such magnetic experiments provided access to an estimate of both the cell elasticity and of the adhesion energy of cell-cell interaction.

This relationship between the macroscopic behavior of a cellular aggregate and the microscopic properties of cell components sheds light on the material properties of cell assemblies and has important implications on the attempts to create functional tissue structures *in vitro*.

#### 2.3. Magnetic endothelialization of tubular scaffolds with EPCs: towards vessel substitutes

Controlled magnetic cellular organization was developed to create structures resembling blood vessels starting from tubular gels. The aim was to create tissue substitutes with optimized geometries and cellular organizations, possibly combining several cell types.

The advantage of magnetic patterning, using magnets placed outside the substrate that supports the cells, is that it is directly applicable to a thick 3D biomaterial. Here, a magnetic force was used to generate, within a porous tube scaffold, a cellular organization approaching that of native vessels. The goal was to obtain a geometry in which EPCs are located on the tube lumen, and smooth muscle cell precursors (MSCs) are located within the tube[67]. For this purpose, a circular permanent magnet with a radial field gradient was used to magnetically attract the EPCs on the lumen, as described in Figure 2D. The tube was first seeded with non-magnetic MSCs then placed at the center of the magnet and magnetic EPCs were injected into the lumen (Figure 4A). It has to be noted that the full compatibility of the magnetic labeling for EPCs was demonstrated, and particularly the preservation of endothelial capillary formation with the magnetic cells[90,94].

**Figure 4:**
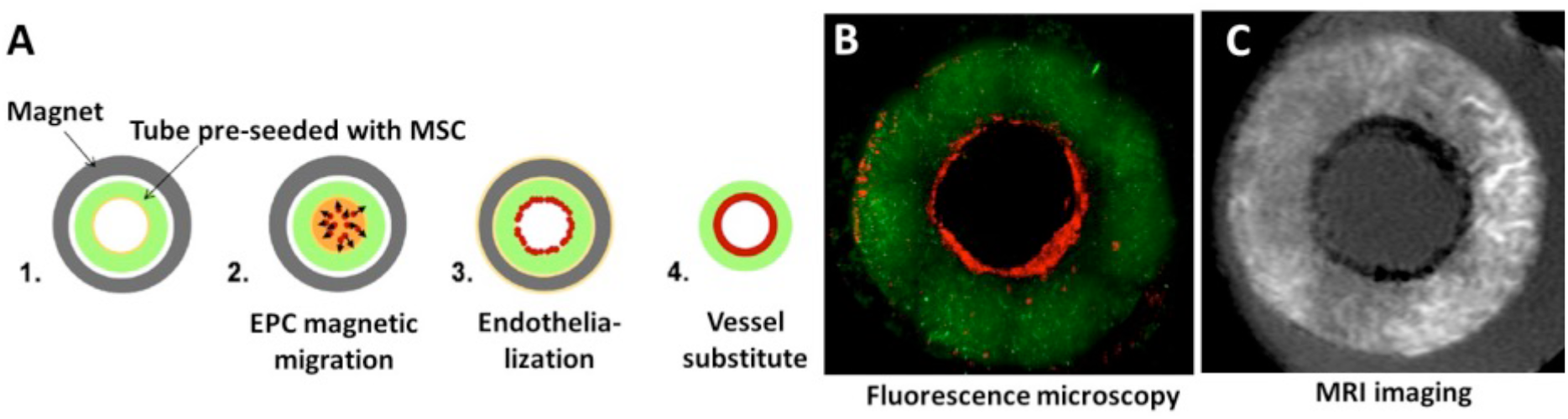
A. To fashion vessel substitutes, tubular gels were seeded with MSCs (labeled for instance with a green PKH67 fluorescent marker) and a circular empty magnet was used to attract magnetically labeled EPCs (labeled with a red PKH26 fluorescent marker) towards the tube lumen. B. The final vessel-like construct is observed by fluorescence microscopy, in green for the MSCs seeded within the core of the tube, in red for the EPCs attached to the lumen by magnetic patterning. C. Non-invasive imaging by MRI of the tube, with MSCs appearing in white due to a gadolinium labeling (T1 agent), and EPCs being detected in black due to the presence of the magnetic nanoparticles (T2 agent). Adapted with permission from Di Corato et al.[95]. Copyright 2013 American Chemical Society.

Vascular tissue engineering still needs the development of small diameter vascular grafts that mimic the native structure and cell composition of vessels. Here, the combination of polysaccharide scaffolds with tubular geometry and a magnetic endothelial patterning allowed creating pluricellular and organized vascular grafts of 2 mm of diameter (Figure 4B).

Down the road in the development of tissue engineered products, imaging methods to control the precise fate of these cell therapy products after their implantation in the body must be developed, combining cellular-scale resolution with the ability to detect several cell types simultaneously at any tissue depth within a living organism. MRI is one of the most interesting biomedical imaging methods, as it provides both morphological and functional information while avoiding the use of ionizing radiation. It was possible to visualize by high-resolution MRI the two stem cells types, MSCs and EPCs, by using distinct gadolinium and iron oxides labelling, respectively[95]. The two cell types remained visible, one in black and the other in white, when embedded in the polysaccharide tubular scaffolds (Figure 4C). This method provides cellular-level resolution, deep within the body, along with the corresponding anatomical image. Having no depth limit, this approach has a key advantage over the few techniques currently available for imaging multicellular tissue, most of which are based on multiphoton microscopy.

#### 2.4 Magnetic confinement of MSCs and magnetic spheroids fusion: towards cartilage tissue engineering

To promote chondrocyte differentiation, in addition to biochemical triggers, MSCs need to be compacted densely one to the other, generally in 3D spheroids. The classical method consists of centrifuging a cell suspension and then adding a differentiation medium (containing growth factors) to the cell pellet (Figure 5A). Using magnetic cell aggregation to pack the MSCs in 3D could help reproduce, or rather improve, the process[36]. In particular, current limitation in cartilage tissue engineering is that only small (submillimetric) spherical spheroids can be delivered (Figure 5B). The number of cells needed for successful chondrogenesis after centrifugation then lies in a very narrow window of around 200 000 cells. The first challenge was to determine if the magnetic compaction could help to overcome this size limitation. This was not the case; they have the same size limit due to limited diffusion of nutriments and growth factors to the spheroid core of large-sized ones.

**Figure 5:**
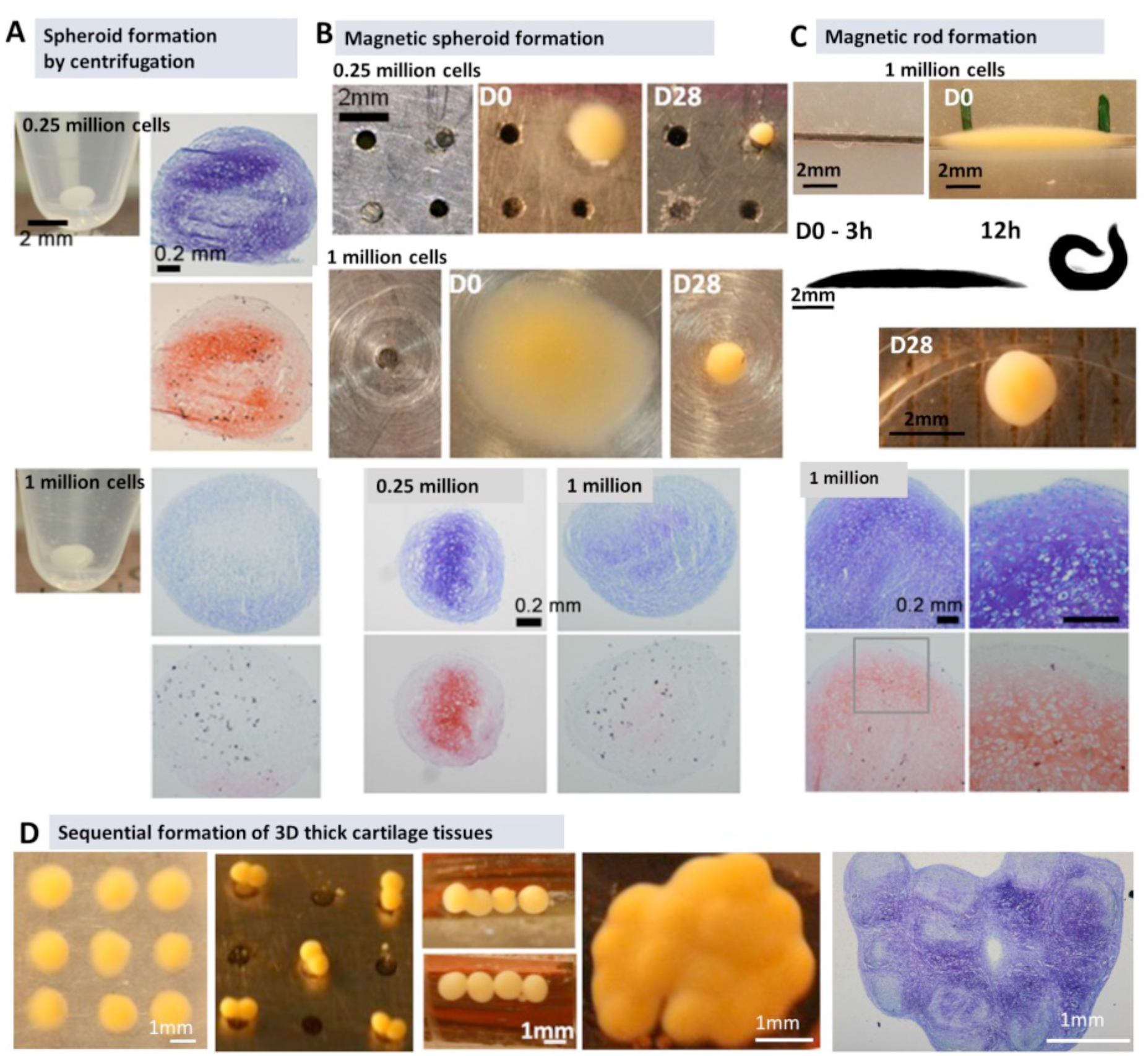
A: Spheroids of 250 000 and 1 million MSCs were formed by centrifugation and then differentiated as cartilage. The 250 000 cells spheroids obtained expressed gycosaminoglycans (typically synthesized in cartilage) after 28 days of chondrogenic differentiation, as demonstrated by the positive toluidine blue (top row) and safranin O (bottom row) stainings. B: Spheroid formation was also performed magnetically, by attracting a given number of stem cells (here 250 000 and 1 million) to a magnetic tip. The cell pellets obtained are more compact when formed magnetically than by centrifugation. Toluidine blue (top row) and safranin O (bottom row) stainings for glycosaminoglycans detection indicate a more abundant production when the spheroids are composed of 250 000 cells. C: Magnetic cell patterning can also be used to create rod-shaped aggregates. In this case, 1 million MSCs are seeded on top of a magnetic line. The linear cell aggregate curves within 12 hours and evolves into a fully formed sphere at day 4-5, which contains glycosaminoglycans after 28 days of chondrogenic differentiation (positive toluidine blue (top row) and safranin O (bottom row) stainings). D: Spheroids of MSCs formed magnetically can be used to create scaffold-free tissues. Independent spheroid units are first created as described previously (via cell-cell contact that leads to a compact structure within a few days). These spheroids can be fused two-by-two then four-by-four up to creating a mm-scale scaffold-free tissue. Cell-cell contact combined with chemical factors allowed to drive the differentiation of the MSCs into chondrocytes and develop a large cartilage tissue containing glycosaminoglycans (toluidine blue staining on the right image). Reproduced with permission from [36].

Next, the shape control elicited by the magnetic patterning was taken advantage of by creating rod shaped tissues (Figure 5C) containing much more cells, but with small lateral diameter, which still allows for nutriment diffusion[96]. Such rod-shaping is clearly impossible for cell pellets prepared by centrifugation. What occurred however was quite unexpected, as a spontaneous shape transition from tissue rods to sphere-like structures happened. However, the final sphere was still overcoming the existing limitations, being four times larger than the current size threshold, and presenting successful chondrocytes differentiation with collagen II and proteoglycans production.

Finally, the MSCs magnetic spheroid patterning were used to create large tissue substitutes of variable geometry with a high potential for future integration[36]. In this case, MSCs spheroids were formed and then fused using magnetic forces. It resulted in the merging of single spheroid units into a bigger tissue structure, of which the final size and shape were magnetically controlled by micromagnets of different geometries (Figure 5D). Thick tissue sheets of functional cartilage tissue were produced with this method. Spheroids made of magnetic MSCs can also be stimulated remotely using a magnetic field. Such exposure increased the chondrogenic differentiation of adipose-derived MSCs, loaded with magnetic nanoparticles, and exposed to a magnetic field every 2 hours for ten minutes during the first 7 days after chondrogenic induction[97].

### 3. The magnetic fingerprint of tissue as a driver of external stimulation

#### 3.1 Compression of magnetic spheroids to retrieve their surface tension

The magnetic fingerprint of the magnetically formed spherical aggregates, or spheroids, conferred by the magnetized cells prior to formation can be used to measure their surface tension, in itself an important marker of tissue cohesion**[98]** and in the case of cancer spheroids, a marker of cell invasion**[99]**. In this case, the stimulation can be performed within 24-72 hours of tissue maturation (without a magnetic force) by subjecting the spheroid to a magnetic field in order to induce a deformation when on a flat surface. The magnetic force required to induce this deformation is equivalent to a magnetic “super-gravity” of around 100 g, quite similar to a centrifugal force. This stimulation forces the spheroid to adopt a compressed shape which reaches equilibrium within minutes. This deformation in super-gravity provides the surface tension parameter of the whole spheroid, mirroring the cells intra-spheroid interactions. Such measurement has been performed not only on stem cells spheroids**[100]** but also on cancer cells spheroids, with a direct implication to measurement of cancer cells invasiveness**[93]**.

#### 3.2 Magneto-mechanical induction of ESCs mesoderm differentiation: towards cardiac tissue engineering

A major challenge in tissue engineering remains to manage, after tissue building, to stimulate mechanically the 3D tissue-like structure at will. It is of relevance for cardiac development, where tissues are continuously subject to pulse contraction.

Cell spheroids composed of ESCs can be formed magnetically following a similar protocol to the one described in the previous section and can serve as initial units for cardiac tissue formation.

Furthermore, magneto-mechanical stimuli can then be applied. Remarkably, such stimulation improved the mesodermal differentiation. In this case, the magnetic micro-attractor used for 3D cell assembly also functions as a micro-controlled attractor network to manipulate the model tissue without direct contact[43]. This approach provides an “all-in-one” solution: the same device is used to create the model tissue and to stimulate it throughout its maturation, for example in a cyclical manner that mimics cardiac muscle contraction. The principle was to form a spheroid of ESCs on one micro-magnet (Figure 6A), then to trap the spheroid by approaching a second similar micro-magnet and stretch at will the maturing embryoid body (Figure 6B) in between. When imposing a cyclic stimulation, magnetic ESCs differentiation was driven towards the cardiac pathway (Figure 6D & 6C). This constitutes the first tried and tested approach allowing the creation of an embryoid body with no direct contact and no supporting matrix, followed by its maturation and stimulation *in situ*.

**Figure 6:**
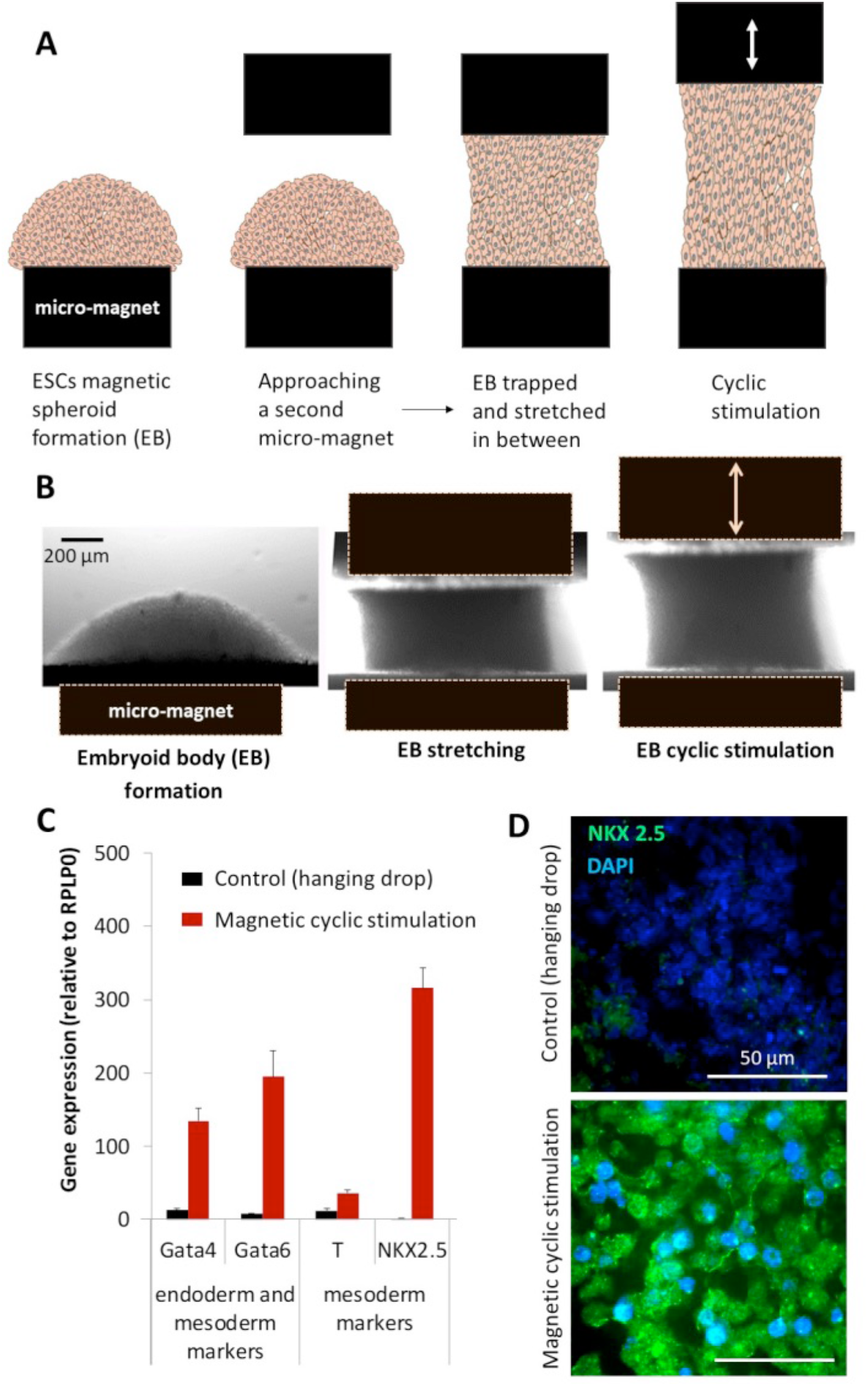
A. Principle of the magnetic tissue builder and stretcher: A first micro-magnet (such as the one described in Figure 2A) aggregates the magnetic ESCs together before a second magnet is used to stretch and stimulate (for instance, cyclically) the created tissue (herein an embryoid body). B. Typical bright field images of embryoid body formation, stretching in between the two micro-magnets after a second one is approached, and cyclic stimulation. C. Expression of genes typically expressed in the mesoderm after 5 days of differentiation was assessed for embryoid bodies formed magnetically and then subjected to cyclic stimulation and compared to embryoids created via the hanging drop method (control). D. Immunofluorescent images were performed on the control and on samples subjected to cyclic stimulation and stained with NKX 2.5 (marker of differentiation, in green) and DAPI (cell nuclei, in blue).

A higher-throughput magnetic micro-array containing 900 micro-magnets served a similar purpose by providing the advantage of creating 900 embryoid bodies (EBs) in one single pipetting step. These EBs could then be compacted cyclically by placing an electromagnet underneath. The cyclic magnetic compaction applied during the first fundamental stages of differentiation increased it toward the cardiac mesoderm pathway, similarly to the differentiation obtained with single EBs, but with high throughput in this case [101].

## Conclusion

Overall, magnetic forces can be powerful tools to organize and differentiate cells, and in particular stem cells in different mechanical microenvironments.

Stem cells were easily confined in a controlled configuration (spheroidal, tubular) thanks to magnetic substrates of variable architecture, at sub-millimetric and millimetric scales. Magnetic forces are applied to the cells remotely and can be modulated at will, in space and/or time. They can also create a specific mechanical environment (such as a cyclic contraction) sufficient to improve *in vitro* tissue functionality.

Importantly, stem cells needed first to be magnetized by allowing them to internalize magnetic nanoparticles. The biocompatibility of these nanoparticles is excellent, and the magnetic labeling generally does not affect cell function. However, before considering their use for regenerative medicine applications, it is crucial to study their long-term intracellular fate within tissues. Stem cell spheroids appear as a well-designed tool to monitor intracellular nanoparticle degradation. Indeed, each single spheroid’s magnetism is a fingerprint of nanoparticles integrity, and therefore of potential degradation. Remarkably, it was found that magnetic nanoparticles can be more than 90% purged inside the stem cells spheroids in the first ten days of tissue maturation[102]. This near-total nanoparticle degradation yet barely affected cellular iron homeostasis, which bodes well for their safety in regenerative medicine applications. Also worth mentioning, the massive biodegradation of the nanoparticles’ magnetic core can be prevented by fine-tuning an inert gold shell[103]. This can be important for long-term use of the nano-functionality, for instance for monitoring the outcome of cell therapy by magnetic resonance imaging, or for applying multiple stimulation on an engineered tissue. It was recently demonstrated that MSCs are capable of biosynthesizing magnetic nanoparticles upon transformation of previously internalized man-made nanoparticles[68,104]. This process opens to the potential of producing stable and fully biogenic nanoparticles that can be 100% biocompatible.

The development of magnetic tools to improve tissue engineering and understand cell differentiation involves cross-disciplinary scientific and technological approaches in microfabrication, physics and life sciences. The multidisciplinary nature of the research achieved combines nanoparticles, magnetic field miniaturization, cellular biology, and tissue engineering. The field of tissue engineering, as it relates to the study of stem cells, is receiving input from the fields of cell biology, chemistry, biomechanics and medicine. With magnetism, it adds a physical approach to cell differentiation and tissue regeneration, placing the focus on the influence of mechanical constraints.

## Acknowledgments

The writing of this review article was supported by the European Union (ERC-2019-CoG project NanoBioMade #865629).

## Notes

### Competing Interest Statement

The authors have declared no competing interest.

